# Social sensing of infection reprograms peripheral immunity in healthy mice

**DOI:** 10.64898/2026.01.28.702380

**Authors:** Temitope W. Ademolue, Lena Pernas

## Abstract

In plants and insects, social immunity enables individuals to detect infection in neighbors and mount protective, community-level responses. Whether mammals possess analogous mechanisms remains unknown. Here, we asked how the presence of sick cage-mates influences the physiology of uninfected neighbors. We found that healthy mice co-housed with conspecifics infected with the non-communicable murine pathogen *Toxoplasma gondii* undergo a shift in peripheral immune responses that establishes a primed immune state. This exposure-induced priming conferred physiological resilience to a sublethal lipopolysaccharide (LPS)-inflammatory challenge and was mediated by increased myeloid-derived IL-10 production. Blocking IL-10 signaling abrogated exposure-induced protection against a subsequent immune challenge. Thus, our findings show that immune state in healthy mammals can be shaped by exposure to infected conspecifics, hinting at social immunity-based protective mechanisms in mammals.

**One sentence summary:** Immune responses in healthy mammals are shaped by exposure to infected conspecifics.

## Introduction

Microbial infection has posed a recurrent and profound threat to human populations, as exemplified by the most recent COVID-19 pandemic ^1^. We have made significant progress in defining how infected mammals mount immune responses to combat pathogens ^2–5^. By contrast, whether infected individuals prepare healthy neighbors against subsequent immune challenges remains unknown. This gap in our knowledge reflects a prevailing focus in mammalian immunology on host-pathogen interactions within the infected individual. Thus, despite well-establish population-level concepts such as herd immunity, our understanding of how immune states may be shaped by social interactions among individuals remains little understood.

Discoveries in other biological systems have demonstrated that immunity extends beyond the individual and enables organisms to detect infection in neighbors and mount anticipatory protective responses. Indeed, infection-induced communication in plants can activate or prime immune defenses in uninfected neighbors ^6,7^. For example, tobacco plants infected with the tobacco mosaic virus (TMV) release methyl salicylic acid (MeSA), a volatile compound that is detected by neighboring plants and leads to acquired resistance to subsequent infections ^8^. Green leaf volatiles released from infected plants can also prime neighboring plants into a heightened state of immune readiness, enabling accelerated defensive responses upon later pathogen exposure ^9^.

Social immunity has also been described in eusocial insects. Nurse honeybees transmit information about environmental pathogens to colony members through the royal jelly they produce, which is consumed by the queen throughout her life and worker-larvae for the first three days of life ^10^. Pathogen-derived molecules ingested by nurses are incorporated into royal jelly and induce antimicrobial responses in the larvae that consume them, thereby conferring protection prior to infection ^11,12^.

In mammals, signals emitted by infected individuals are known to alter the behavior of healthy cage-mates ^13^. Olfactory cues derived from the excretions of virally infected mice elicits avoidance behaviors in uninfected neighbors ^13,14^. Visual cues transmitted by infectious avatars lead to changes in peripheral immune responses in humans exposed through virtual reality ^15^. However, whether sensing infection in a conspecific can alter immune function in neighboring healthy mammals—and thereby influence their response to subsequent immune challenges—remains unknown.

Here, we asked whether mammals detect infection in neighboring cage mates, and whether such detection influences physiological responses in healthy individuals. We found that exposure to mice infected with the non-communicable pathogen *Toxoplasma gondii* led to increases in anti-inflammatory cytokines including IL-10 in healthy mice and promoted their resilience to a subsequent inflammatory challenge. Blocking IL-10 signaling abolished this exposure-induced protection. These findings demonstrate that exposure to infected cage mates can alter immune responses in healthy mammals, revealing a previously unrecognized form of social immunity in mammals.

## Results

### Exposure to infected animals induces resilience to a sublethal LPS challenge

To determine whether exposure to an infected mouse affected subsequent responses to inflammatory challenges, we chose as an infection model the pathogen *Toxoplasma gondii* for two reasons. First, rodents are intermediate hosts of *Toxoplasma* and infection is therefore non-communicable between conspecifics ^16^. Second, acute toxoplasmosis in mice infected with Type I *Toxoplasma* parasites manifests in an early, asymptomatic phase, followed by a symptomatic phase characterized by rapid weight loss, morbidity, and inflammation ^17^. We confirmed that up to 4 days post infection (dpi), mice maintained stable core body temperature, body weight, and glucose levels (**Figure 1A-B**). After 4 dpi however, mice exhibited progressive weight loss, hypothermia, and elevated ketone bodies and circulating inflammatory cytokines (**Figure 1A-B, S1A**) ^17,18^.

**Figure 1:**
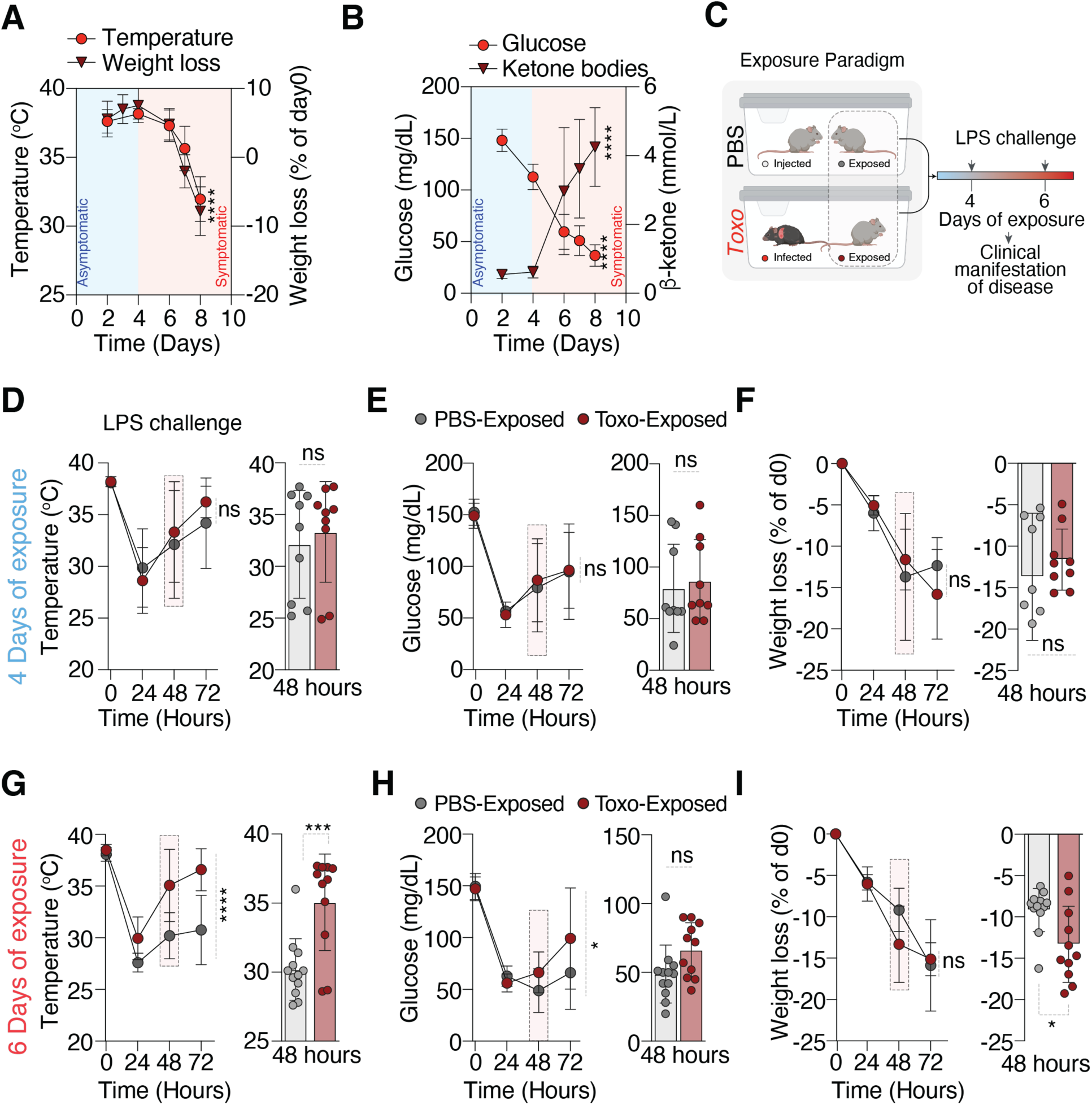
Exposure to infected animals enhances physiological resilience to inflammatory stress. **(A)** Core body temperature (red circles) and body weight loss (dark red triangles) measured at indicated days post infection (dpi) in *Toxo*-infected animals, shown across asymptomatic (blue shading) and symptomatic (red shading) phases. Data are mean ± SD, \*\*\*\**P* <0.0001 by two-way ANOVA (row factor, corresponding to time effect) (*n* = 6 mice per group). **(B)** Blood glucose (red circles) and β-ketone body levels (dark red triangles) measured at indicated dpi in *Toxo*-infected animals. Data are mean ± SD, \*\*\*\**P* <0.0001 by two-way ANOVA (*n* = 3-6 mice per group). **(C)** Schematic of the exposure paradigm. Mice were exposed to PBS-injected or *Toxo-*infected cage mates for 4 or 6 days, followed by 2.5 mg/kg LPS challenge. **(D)** Core body temperature, **(E)** blood glucose and **(F)** body weight loss following LPS challenge in mice exposed to PBS-injected and *Toxo-*infected cage mages for 4 days. Line plots show longitudinal measurements; bar plots show values at 48 hours after challenge. Data are mean ± SD, ns, not significant by two-way ANOVA (exposure) and unpaired t-test for timepoint pairwise comparison (*n* = 12 mice per group). **(G)** Core body temperature, **(H)** blood glucose, and **(I)** body weight loss following LPS challenge in mice exposed to PBS-injected and *Toxo-*infected cage mages for 6 days. Line plots show longitudinal measurements; bar plots show values at 48 hours after challenge. ns, not significant; data are mean ± SD, \**P* < 0.05; \*\**P* < 0.01; \*\*\**P* < 0.001; \*\*\*\**P* < 0.0001 by two-way ANOVA and unpaired t-test for timepoint pairwise comparison (*n* = 9-12 mice per group). Data in D-I are pooled from two independent experiments.

We next leveraged these distinct phases of *Toxoplasma* infection to establish an exposure paradigm. To do so, three naïve mice were co-housed with either three PBS-injected or three *Toxoplasma*–infected cage mates, which we will hereafter refer to as PBS-exposed and *Toxo-*exposed, for the asymptomatic phase (4 dpi) or the duration of both the asymptomatic and symptomatic phases (6 dpi) (**Figure 1C**). During the exposure period, animals remained in the same home cage with minimal handling stress. The healthy PBS- and *Toxo-*exposed mice were subsequently challenged with a sublethal dose of lipopolysaccharide (LPS), a potent inducer of systemic inflammation, and monitored for 3 days ^19–21^. Mice exposed to both PBS-injected and *Toxo*-infected cage-mates for 4 days exhibited comparable decreases in core body temperature, blood glucose and body weight at 24 hpi, with partial recovery by 72 hours as expected (**Figure 1D-F**) ^19–21^. By contrast, following 6 days of exposure, *Toxo-*exposed mice exhibited a marked improvement in recovery of core body temperature and blood glucose beginning at 48 hours post LPS-challenge relative to PBS-exposed controls (**Figure 1G-H**). This advantage was maintained throughout the 72-hour observation period and was accompanied by greater weight loss in *Toxo-*exposed mice (**Figure 1G-I**). Thus, cumulative exposure to the acute asymptomatic and symptomatic phases of *Toxo*-infected animals promotes thermoregulation and glucose homeostasis during a subsequent inflammatory challenge.

### Exposure to infected animals induces an IL-10 signaling program

How does exposure to *Toxo-*infected cage-mates promote resilience to a sublethal LPS challenge? To exclude the possibility that this effect resulted from parasite transmission—although not expected between intermediate hosts—we compared transcript levels of the *Toxoplasma* major surface antigen (*Sag1*) gene between *Toxo*-infected and *Toxo*-exposed mice. At 4 and 7 dpi, *Sag1* transcripts were readily detectable in infected mice in liver and small intestine—sites of highest *Toxoplasma* burden following an intraperitoneal infection—but not in *Toxo*-exposed mice (**Figure 2A, S1B**) ^22^. Consistent with the absence of infection, core body temperature and blood glucose levels in *Toxo*-exposed mice were similar to PBS-exposed controls during the exposure period (**Figure S1C**).

**Figure 2:**
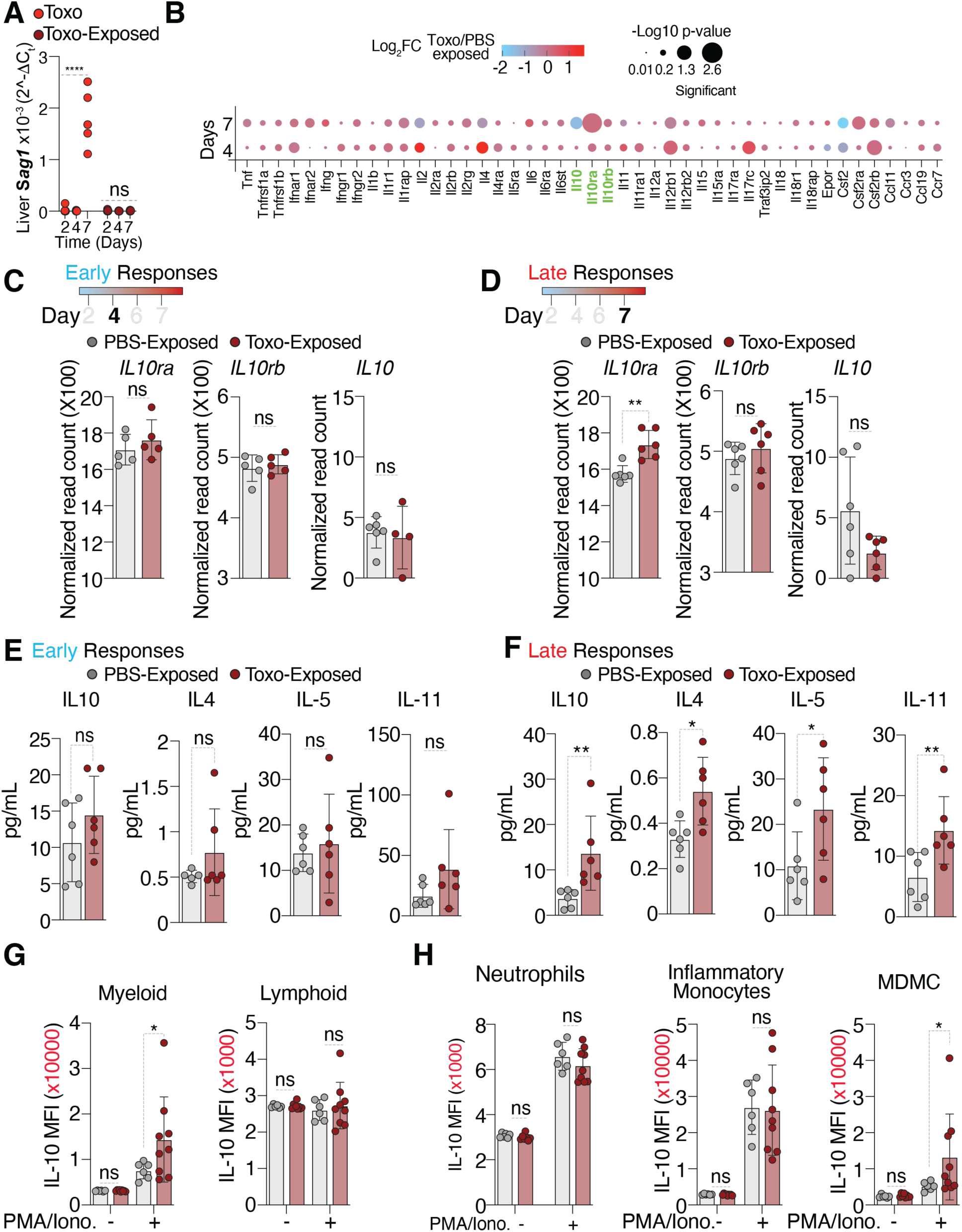
Exposure to infected animals induces an IL-10 signaling program. **(A)** Liver *Sag1* transcript abundance measured at 2, 4, and 7 dpi in infected (*Toxo*) and 2, 4, and 7 days post exposure to *Toxo-*infected animals. ns, \*\*\*\**P* < 0.0001 by two-way ANOVA comparing day 2 versus day 7 within *Toxo* and within *Toxo*-exposed groups (*n* = 3-6 mice per group). **(B)** Bubble plot showing differential expression of cytokine and cytokine receptor genes in spleen from RNAseq analysis of *Toxo*-exposed relative to PBS-exposed mice at 4 and 7 days post exposure. Color indicates log_2_fold change (*Toxo*/PBS exposed) and bubble size reflects −log_10_ *p*-value, *n* = 4-6 mice per group from 2 independent experiments. **(C)** Normalized read counts for *Il10*, *Il10ra*, and *Il10rb* from spleens as in (B) at days 4 and **(D)** 7. ns, not significant; \*\**P* < 0.001 by unpaired t test; data are mean ± SD for *n* = 4-6 mice per group. Data are pooled from two independent experiments. **(E)** Serum cytokine concentrations at days 4 and **(F)** 7 for IL-10 IL-4, IL-5 and IL-11. Data are mean ± SD for *n* = 6 mice per group from two independent experiments; ns, not significant; \**P* < 0.05; \*\**P* < 0.01 by unpaired *t*-test. **(G)** IL-10 median fluorescence intensity (MFI) in myeloid and lymphoid populations isolated from peripheral blood from 7 day-PBS-exposed and *Toxo*-exposed mice following *ex vivo* stimulation with or without 1X PMA/ionomycin. ns, not significant. Data are mean ± SD, \**P* < 0.05 by two-way ANOVA for *n* = 6 per PBS-exposed group and 9 mice per *Toxo-*exposed group. **(H)** IL-10 MFI in neutrophils, inflammatory monocytes and monocyte-derived macrophage-like cells (MDMCs) from PBS-exposed and *Toxo*-exposed mice with or without PMA/ionomycin stimulation. Data are mean ± SD, ns, not significant; * *P* < 0.05 by two-way ANOVA for *n* = 6 per PBS-exposed group and 9 mice per *Toxo-*exposed group.

We next asked whether, analogous to priming in plants, exposure to infected conspecifics induced a primed state that underlies the improved temperature and glucose recovery following LPS challenge ^9^. We reasoned that such a primed state may result from systemic metabolic or immune alterations in *Toxo-*exposed animals. To address this possibility, we first performed untargeted metabolomics of serum samples from PBS-exposed and *Toxo-*exposed mice after exposure limited to the asymptomatic phase (4 dpi), or after cumulative exposure to the acute asymptomatic and symptomatic phases (7 dpi), the latter of which conferred resilience during LPS challenge (**Figure 1D-I**). However, principal component analysis (PCA) revealed no separation between PBS-exposed and *Toxo-*exposed groups at either time point (**Figure S1D-E**). Furthermore, measured metabolites were similar at either day 4 or day 7 after exposure (**Figure S1D-E**).

To determine whether immune function was altered in *Toxo*-exposed animals, we first examined the spleen, a secondary lymphoid organ that senses blood-borne inflammatory signals and coordinates innate immune responses ^23^. We compared the transcriptome of spleens isolated from PBS-exposed and *Toxo*-exposed mice after 4 and 7 days of exposure. Unsupervised PCA revealed no global transcriptional separation between *Toxo*-exposed and PBS-exposed mice at all time points examined (**Figure S1F**). However, targeted analysis of a panel of immune-related transcripts revealed modest yet broad changes after 7 days exposure (**Figure 2B**). Of interest to us were transcripts encoding the ligand-binding α subunit of the IL-10 receptor *(Il10ra)* were significantly increased in *Toxo-*exposed mice (**Figure 2B-D**). IL-10RA specifically mediates signaling by IL-10, an anti-inflammatory cytokine known to attenuate LPS-induced pro-inflammatory responses ^24–26^. By contrast, transcripts of the shared receptor subunit *IL10rb* was unchanged, or of classical inflammatory and effector cytokine receptors including *Il17ra*, *Il17rc*, *Il15ra*, *Il11ra1*, *Il6ra*, *Il5ra*, *Il4ra*, *Il2ra*, *Il2rb*, *Il2rg*, *Il1r1*, *Ifngr1*, *Ifngr2*, *Ifnar1*, *Ifnar2*, *Tnfrsf1a*, and *Tnfrsf1b* were expressed at comparable levels between PBS- and *Toxo-*exposed mice following both 4 and 7 days of exposure (**Figure 2E-F**).

Building on these findings, we next assessed whether the observed transcriptional changes were accompanied by change in systemic immune parameters. To do so, we quantified 44 cytokines, chemokines, and growth factors in the peripheral blood of PBS-exposed and Toxo-exposed mice. After 7 days of exposure, *Toxo*-exposed mice exhibited significantly elevated levels of IL-10, and cytokines associated with tissue tolerance and repair including IL-4 and IL-5 (**Figure 2E-F; S2; Table S1-2**) ^4,27^. 4 days of exposure to *Toxo-*infected mice was not sufficient to induce significant changes in circulating IL-10, IL-4, IL-5, and IL-11 in healthy animals (**Figure 2F; S2A, Tables S1-2**). Thus, exposure to both the asymptomatic and symptomatic stages of *Toxo*plasma infection in cage-mates induced an IL-10-associated cytokine response in healthy mice.

### Exposure to sick animals enhances IL-10 production in peripheral myeloid cells

What drives the increase in circulating IL-10 observed in exposed mice? To begin to address this question, we asked whether exposure to *Toxoplasma-*infected cage-mates altered the abundances of IL-10-producing immune cells in circulation. Because IL-10 can be produced by multiple immune cell types, including helper T cells, regulatory T cells, CD8 T cells, B cells, macrophages and NK cells, we first compared the abundance of CD11b^+^ myeloid cells, and CD11b^−^ lymphoid cells in peripheral blood ^25^. No significant differences in the frequencies of these populations were detected between PBS-exposed and *Toxo-*exposed mice after 7 days of exposure (**Figure S3A-B**). Neither in frequencies of total monocytes, T cells and B cells (**Figure S3C**).

We next compared levels of IL-10 in circulating immune cells with and without stimulation *ex vivo*. Following stimulation, myeloid-derived IL-10 was significantly higher in *Toxo*-exposed mice than PBS-exposed mice, whereas lymphoid-derived IL-10 remained comparable (**Figure 2G; S3A)**. To determine whether this difference was due to a global increase in myeloid-IL-10 production, we compared IL-10 levels across CD11b+ myeloid populations, including inflammatory monocytes (Ly6C^hi^Ly6G^neg^), neutrophils (Ly6C^neg^Ly6G^pos^), and myeloid-derived monocytes (MDMCs), which encompass patrolling monocytes (Ly6C^mid^Ly6G^neg^) ^28^. We found that MDMCs from *Toxo*-exposed mice exhibited significantly higher inducible IL-10 production following stimulation compared to MDMCs from PBS-exposed mice, while baseline IL-10 levels remained unchanged (**Figure 2H**). IL-10 production in inflammatory monocytes and neutrophils however was similar between PBS-exposed and *Toxo-*exposed mice (**Figure 2H**). Thus, enhanced inducible IL-10 production following exposure to *Toxoplasma-*infected cage-mates was restricted to MDMCs.

### Blocking IL-10 abolishes exposure-dependent protection during acute inflammation

Having established that exposure to infected cage-mates modulates IL-10 signaling, we next asked whether IL-10 was required for the exposure-induced physiological resilience to LPS challenge (**Figure 1G-I)**. To address this, PBS-exposed and *Toxo*-exposed mice following 7 days of exposure were treated with either a neutralizing anti–IL-10 antibody or isotype control prior to lipopolysaccharide (LPS) challenge (**Figure 3A**). Core body temperature, glycemia and body weight were monitored over a 72-hour period. Neutralization of IL-10 abolished the improved recovery of core body temperature and glucose levels in *Toxo*-exposed mice relative to PBS-exposed mice after LPS challenge (**Figure 3B-C**). LPS-induced weight loss was unaffected by treatment with anti-IL-10 antibody *Toxo-*exposed mice (**Figure S3D-E**). Of note, administration of anti–IL-10 antibody did not exacerbate the physiological response to LPS in PBS-exposed control mice as core body temperature remained comparable between isotype- and anti-IL10-treated groups (**Figure S3F**). Thus, IL-10 signaling is required for the exposure-induced enhancement of thermometabolic recovery during systemic inflammation (**Figure 4**).

**Figure 3:**
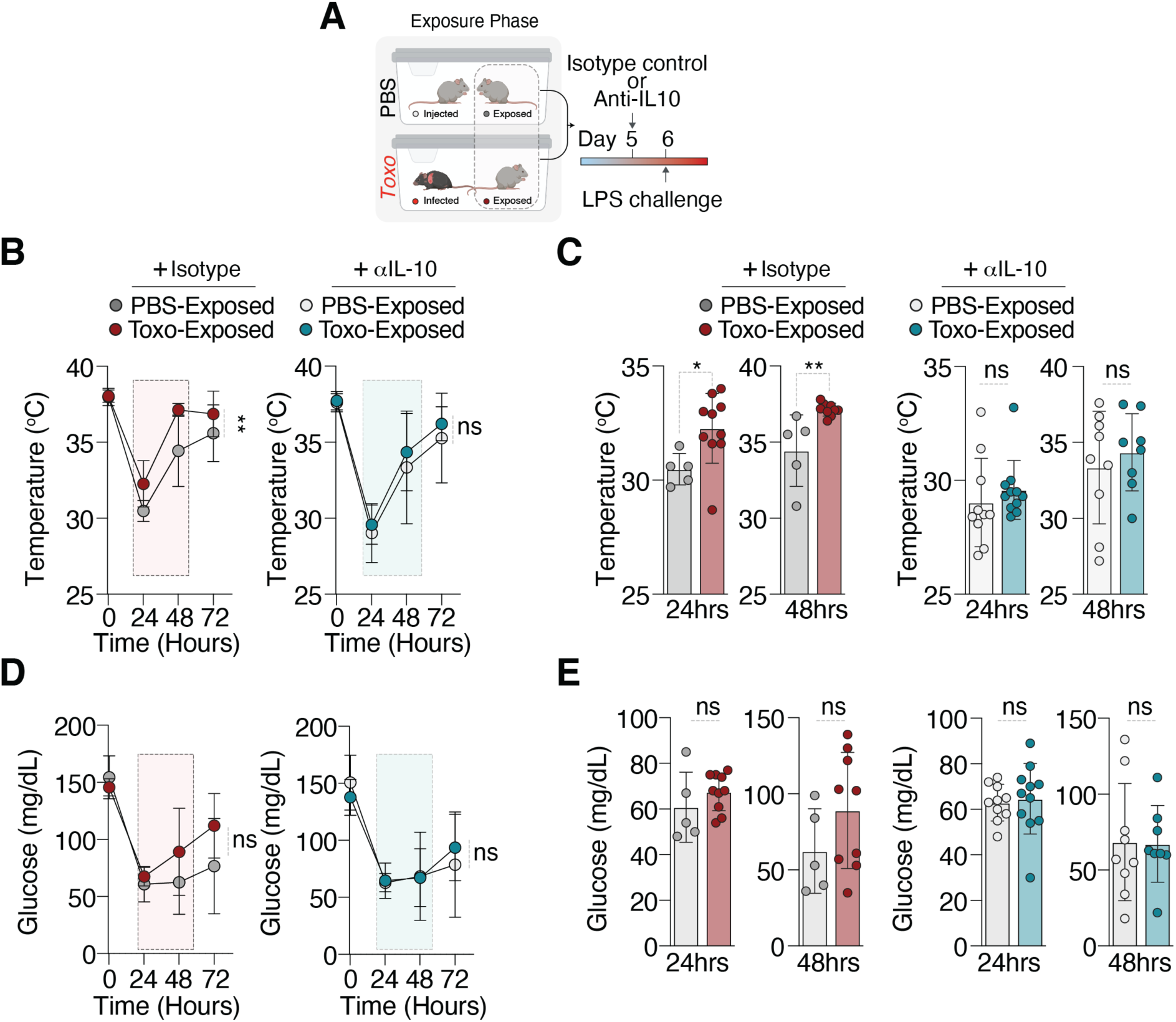
Blocking IL-10 abolishes exposure-dependent protection during acute inflammation. **(A)** Schematic of IL-10 neutralization experiment. At day 5 of the exposure phase, PBS-exposed and *Toxo*-exposed mice received isotype control or anti–IL-10 antibody, followed by LPS challenge (2.5mg/kg) on day 6. **(B)** Core body temperature at indicated hours post injected (hpi) following LPS challenge in isotype-treated (left) or anti–IL-10–treated (right) PBS-exposed and *Toxo*-exposed mice. Data are mean ± SD, \*\**P* < 0.01 by two-way ANOVA. **(C)** Core body temperatures at 24 (Isotype, *n* = 5 PBS-exposed, 10 *Toxo*-exposed; anti—IL-10, *n* = 10 PBS-exposed; 11 *Toxo*-exposed) and 48 hours (Isotype, *n* = 5 PBS-exposed, 9 *Toxo*-exposed; anti—IL-10, *n* = 9 PBS-exposed; 8 *Toxo*-exposed) after challenge of mice as in B. Data are mean ± SD, ns, \**P* < 0.05, \*\**P* < 0.01 by unpaired t-test. **(D)** Blood glucose in mice at indicated hpi following LPS challenge in isotype-treated (left) or anti-IL-10 treated (right) PBS-exposed and *Toxo*-exposed mice. Data are mean ± SD, ns by two-way ANOVA. **(E)** Blood glucose of mice from (E) (Isotype, *n* = 5 PBS-exposed, 10 *Toxo*-exposed; anti—IL-10, *n* = 10 PBS-exposed; 11 *Toxo*-exposed) and 48 hours (Isotype, *n* = 5 PBS-exposed, 9 *Toxo*-exposed; anti—IL-10, *n* = 9 PBS-exposed; 8 *Toxo*-exposed. Data are mean ± SD analyzed by unpaired t test. Data pooled from two independents. Only animals that met a predefined physiological response criterion—development of hypothermia (<35 °C) within 24 hours of LPS injection—were included in the analysis.

**Figure 4:**
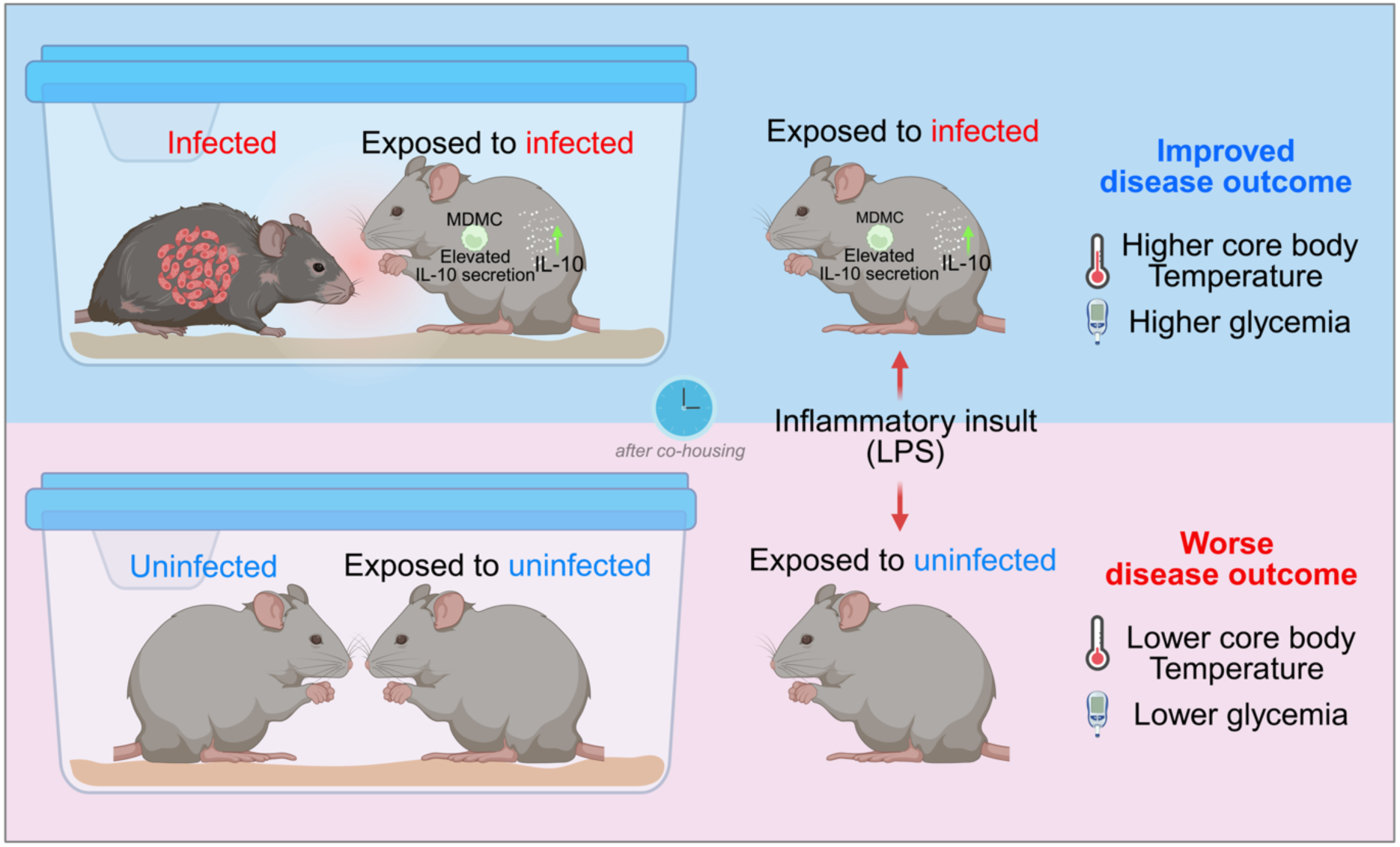
Social sensing of infection reprograms peripheral immunity in healthy mice. Exposure to *Toxo*-infected mice induces enhanced IL-10 production from peripheral myeloid derived monocyte cells (MDMCs) in healthy neighboring mice. Upon a subsequent inflammatory LPS challenge, mice exposed to *Toxo*-infected mice have higher core body temperature and glycemia and thus improved disease outcome. Created with Biorender.com.

### Exposure-induced thermoregulatory resilience extends to live *Toxoplasma* infection

We next tested whether exposure to infected cage-mates altered physiological outcomes during a pathogen infection. Naïve mice were exposed for either 4 or 6 days to PBS-injected or *Toxoplasma*–infected cage-mates and subsequently challenged with 50 *Toxoplasma* tachyzoites (**Figure S4A**). Following *Toxoplasma-*injection after 4 days of exposure, both PBS-exposed and *Toxo*-exposed mice developed hypothermia during infection (**Figure S4B**). However, at 8 dpi—immediately prior to the humane endpoint and corresponding to peak disease severity—*Toxo*-exposed mice maintained significantly higher core body temperatures than PBS-exposed controls (**Figure S4B**). In contrast, blood glucose levels declined similarly in both groups (**Figure S4C**). Thus, exposure to the asymptomatic phase of *Toxoplasma-*infection conferred a slight thermoregulatory advantage during a subsequent lethal microbial challenge. Unexpectedly, when infection was initiated after 6 days of exposure, this thermoregulatory advantage was no longer observed (**Figure S4D-E**).

To test whether this effect was generalizable across parasite strains with distinct virulence properties, we repeated the exposure-and-infection challenge paradigm using the Δmyr1 *Toxoplasma* strain, which is defective in host cell effector export and exhibits reduced virulence ^29^. Consistent with our findings using WT parasites, mice exposed for 4 days to infected cage-mates maintained higher core body temperatures than PBS-exposed controls at 8 dpi, despite similar decreases in blood glucose (**Figure S4F-G**). Consistently, this thermoregulatory difference was absent when infection was initiated after 6 days of exposure (**Figure S4F-G).** Thus, exposure to asymptomatic infected cage-mates enhances thermoregulatory resilience during subsequent pathogen challenge.

## Discussion

Our data demonstrate that exposure to infected mammals primes immune responses in healthy conspecifics and raises a fundamental question: how is infection in one individual sensed and translated into immune modulation in another (**Figure 4**)?

A growing body of work highlights bidirectional crosstalk between the nervous and immune systems, whereby perceived environmental signals can shape both innate and adaptive immunity ^30–32^. For example, the activation of the hypothalamic-pituitary-adrenal axis regulates lymphocytes and monocytes migration to the bone marrow during acute stress ^33^. In this framework, sensory cues emitted by infected individuals at different stages of disease progression may exert an influence on the immune state of neighboring animals through neuroimmune signaling pathways.

Multiple, non-exclusive mechanisms may underlie this social transmission of immune state. At early stages of infection, volatile organic compounds (VOCs) which are known to change during infection, may represent a primary mode of communication between infected animals and their neighbors ^34^. At later stages, perceptible changes in behavior or appearance, such as reduced mobility or weight loss, could provide additional sensory cues. Alternatively, immunomodulatory signals may be transmitted through contact-mediated routes, including coprophagy, which is common in rodents ^35^.

Notably, the physiological consequences of exposure depended on both the duration of exposure and nature of the subsequent challenge. Exposure to symptomatic, 6 day-infected cage-mates enhanced resilience to an LPS-induced inflammatory challenge. Yet, this same exposure failed to confer any physiological benefit—and in some contexts diminished—during a subsequent *Toxoplasma* infection. One interpretation is that induction of IL-10, a potent regulator of anti-inflammatory immune programs, is beneficial in settings that require dampening of excessive inflammation, such as endotoxemia, but may be maladaptive during severe microbial challenge where robust antimicrobial responses are required ^26^. This trade-off mirrors observations in plants, where inducible resistance pathways enhance protection against specific pathogens, but can simultaneously increase susceptibility to other ecological threats such as herbivory ^7^. Our findings raise the possibility that socially acquired immune states may confer protection in some contexts while imposing costs in others.

Whether analogous mechanisms operate in humans remains an open but intriguing question. VOCs are powerful regulators of behavior and physiology across vertebrates. During illness, humans emit body odors that evoke avoidance or disgust in healthy individuals ^36^. Odor cues from humans undergoing transient inflammatory challenges are perceived as more aversive, suggesting that immune state can be externally signaled and socially perceived. More recently, exposure to virtually infected human avatars was shown to alter connectivity patterns between infection-sensing brain regions and the hypothalamus, and modulate frequency and activation of innate lymphoid cells in the periphery ^15^. These observations are consistent with the possibility that infection-associated sensory cues can influence immune function in humans, even in the absence of direct pathogen exposure.

Together, our work shows that exposure to infected mammalian conspecifics can modulate immune state in healthy neighbors and influence physiological responses to subsequent immune challenges. These findings reveal a previously unrecognized form of social immunity in mammals, whereby infection-associated cues regulate immune function beyond the infected individual. More broadly, they raise the possibility that socially transmitted signals could be leveraged to promote resilience to inflammatory stress at the population level—while also highlighting the context-dependent trade-offs inherent to socially acquired immune states.

## Acknowledgments

We thank Dr. Joanne Zahorsky-Reeves (Division of Laboratory Animal Medicine (UCLA) for support with animal protocols, UCLA Technology Center for Genomics & Bioinformatics (TCGB), UCLA Metabolomics Core, and General Metabolics for sample processing. We thank Yanying Dai for technical help with mouse experiments, we thank Drs. Sophie Stecolorum, Carlos Ardanaz, Alexander Hoffmann, and Timothy O’Sullivan for insightful discussions on experimental design and interpretation; Hatoon Baazim and Tim Bartsch for helpful and critical manuscript feedback, and all members of the Pernas laboratory for thoughtful discussions.

## Author contributions

Conceptualization: TWA and LP; Methodology: TWA and LP Investigation: TWA; Funding acquisition: LP; Writing – original draft: TWA and LP; Writing – review & editing: TWA and LP; Project administration and supervision: LP

## Data and materials availability

All data are available in the main text or the supplementary materials.

## Mice

Female C57BL/6J mice (The Jackson Laboratory; JAX stock no. 000664) were obtained at 6-8 weeks of age. Mice were housed at 6 animals per cage under standard specific pathogen-free conditions with ad libitum access to food and water. Animals were acclimatized for at least 48 hours after arrival before being subjected to experimentation. All animal experiments were conducted in accordance with institutional guidelines for the care and use of laboratory animals and were approved by the Chancellor’s Animal Research Committee (ARC) at the University of California, Los Angeles, under protocol ARC-2024-007. All procedures were performed in compliance with applicable local, state, and federal regulations, and every effort was made to minimize animal suffering and to reduce the number of animals used.

### Parasites culture and mouse infection

*Toxoplasma gondii* parasites were maintained by serial passage in human foreskin fibroblast (HFF) monolayers cultured in complete DMEM (cDMEM; DMEM supplemented with 10% heat-inactivated fetal bovine serum). Parasites used in this study were derived from the Type I RH background. For in vivo exposure and infection experiments, the RH strain was used as the infectious agent for *Toxo*-infected mice. For preparation of infectious inocula, HFF monolayers were infected with freshly egressed tachyzoites and incubated for approximately 24 hours, until large parasite-containing vacuoles were readily visible and the monolayer appeared close to lysis. Infected monolayers were then scraped and syringe-lysed to release intracellular parasites. Lysates were clarified by centrifugation at 1500 rpm, the parasite-containing pellet was resuspended in phosphate-buffered saline (PBS), and the suspension was passed through a sterile filter. Tachyzoites were then counted, and serial dilutions were prepared such that a final volume of 200 µl PBS contained 50 tachyzoites. Mice assigned to the *Toxo*-infected group were injected intraperitoneally with 50 tachyzoites in 200 µl PBS. PBS-exposed control animals received a mock inoculum consisting of 200 µl PBS prepared from uninfected HFF cultures that were processed in parallel and subjected to the same scraping, syringe lysis, centrifugation, resuspension, filtration, and dilution steps as infected cultures, thereby controlling for host cell–derived material and handling procedures. Animals were monitored daily and euthanized upon reaching predefined ARC humane endpoints, in accordance with institutional guidelines.

### LPS challenge

Lipopolysaccharide (LPS) was administered by intraperitoneal injection at a dose of 2.5 mg/kg body weight. LPS reagents used in this study were obtained from Cell Signaling Technology (cat. no. 14011S) and MedChemExpress (cat. no. HY-D1056). For all experiments, LPS was prepared fresh in sterile phosphate-buffered saline (PBS) immediately prior to injection. We consistently observed that a subset of animals did not exhibit a physiological response at this dose, defined by the absence of hypothermia ^37^. Therefore, analyses of LPS-induced responses were restricted to animals that developed hypothermia, operationally defined as a core body temperature below 35°C at 24 hours after LPS injection. Animals that did not meet this criterion were excluded from downstream analyses. This approach ensured that comparisons were performed among animals that experienced a comparable inflammatory challenge. Furthermore, although this dose is generally considered sublethal, sporadic mortality was observed in a subset of animals.

### Exposure paradigm

All exposure experiments were performed in standard static cages that were not housed on ventilated rack systems; thus, airflow was not continuous. For exposure to infected cagema tes, mice were co-housed in groups of six animals per cage, consisting of three *Toxoplasma gondii*–infected mice (“*Toxo*-infected”) and three uninfected cage mates (“*Toxo*-exposed”). Control cages contained three mice injected with PBS and housed together under identical conditions. Following establishment of the exposure paradigm, cages were not opened or disturbed except at the predefined experimental intervention time points. Animals remained continuously co-housed for the duration of the exposure period (4 or 6 days, as indicated for each experiment), after which downstream interventions (LPS challenge or live parasite infection) and sample collection were performed. This experimental design ensured prolonged, uninterrupted exposure of naïve animals to infected cage mates while controlling for cage density, handling frequency, and environmental conditions across experimental groups.

### Disease parameters

Disease progression was assessed by longitudinal measurement of core body temperature, blood glucose, and body weight. Core body temperature was measured using a digital rodent thermometer (TK-8851) equipped with a rectal probe for mice (3/4″ length, 0.028″ diameter, 0.065″ tip; RET-3). Blood glucose levels were measured from peripheral blood using an Accu-Chek Performa blood glucose meter (Roche) with Accu-Chek Performa test strips (50-count; Roche, Cat. no. **64540110**). Body weight was recorded using an Ohaus® Compass scale (model **CX221**, 220 g capacity, 0.1 g resolution; Catalog no. **470313-236**). All measurements were performed at the indicated time points by the same operators using identical equipment across experimental groups to minimize technical variability.

### RNA extraction and quantitative RT–PCR

Mice were sacrificed at the indicated time points and perfused in toto with 10 mL ice-cold phosphate-buffered saline (PBS). Spleens were harvested, snap-frozen in liquid nitrogen, and stored at −80 °C until processing. Total RNA was extracted from spleen tissue using a guanidinium–thiocyanate–phenol–chloroform–based method followed by column-based RNA cleanup (New England Biolabs, Cat #: E2050S), according to the manufacturers’ instructions. RNA concentration and purity were assessed prior to downstream applications. Complementary DNA (cDNA) was synthesized using the Transcriptor First Strand cDNA Synthesis Kit (Roche) following the manufacturers’ protocols. Quantitative PCR was performed using SYBR Green chemistry (iTaq Universal SYBR Green Supermix, Bio-Rad) on a QuantStudio™ 7 Flex Real-Time PCR System (Applied Biosystems). Relative gene expression was calculated with Acidic ribosomal phosphoprotein P0 (*Arbp0; Forward: 5’-CTTTGGGCATCACCACGAA-3′ Reverse: 5’-GCTGGCTCCCACCTTGTCT-3’*; *Toxoplasma gondii Sag1;* Forward: 5’-*CTGATGTCGTTCTTGCGATGTGGC*-3’ Reverse:5’ *GTGAAGTGGTTCTCCGTCGGTGT*-3’).

### Serum cytokine measurements

Mice were sacrificed at the indicated time points and whole blood was collected. Plasma was isolated and cytokine concentrations were quantified by Eve Technologies (Canada; (Canada; https://www.evetechnologies.com/) using multiplex cytokine analysis, according to the provider’s standard protocols.

### Bulk RNA sequencing and analysis

Bulk RNA sequencing was performed with support from the UCLA Technology Center for Genomics & Bioinformatics (TCGB). Snap-frozen mouse spleen samples were submitted to the TCGB core facility for bulk RNA-seq library construction and sequencing. Bulk RNA-seq libraries were prepared by TCGB using standard mRNA sequencing workflows (poly(A) selection). Libraries were sequenced on an Illumina NovaSeq X Plus platform using paired-end 2 × 50 bp sequencing, with a target depth of ∼30 million reads per sample. Base calling and demultiplexing were performed by the core facility. Sample-level quality control was performed by examining normalized count distributions. Samples exhibiting aberrant signal dominated by a small number of features, assessed using the ratio of maximum to mean normalized counts (Max/Mean ratio), were excluded from downstream analyses. All subsequent analyses were conducted on the filtered dataset. For visualization, gene sets were extracted and effect sizes (log_2_ fold change) and statistical significance (Wald test p-values) were obtained for each gene–timepoint comparison. Outliers were identified and excluded using an interquartile range (IQR)-based approach (1.5X IQR) prior to visualization.

### Plasma metabolomics

Mouse plasma samples were collected at the indicated time points and stored at −80 °C until analysis. Untargeted metabolomics profiling was performed by General Metabolics. A total of 60 plasma samples were processed according to their established extraction and analysis workflows. Plasma samples were thawed on ice and extracted using a methanol-based protocol to solubilize small-molecule metabolites and remove precipitated proteins. Extracts were further diluted 1:50 in chilled 80% methanol in water (v/v) prior to analysis. A pooled study sample (pSS), generated from aliquots of all study samples, was injected periodically throughout the analytical run to assess technical reproducibility. Metabolite profiling was performed using a flow-injection analysis mass spectrometry (FIA-MS) platform operated in negative ionization mode. Data were acquired on an Agilent 6550 iFunnel LC-MS Q-TOF mass spectrometer coupled to an Agilent 1260 Infinity II quaternary pump and a Gerstel MPS3 autosampler. The running buffer consisted of 60% isopropanol in water (v/v) buffered with 1 mM ammonium fluoride. Mass spectra were acquired in 4 GHz high-resolution mode over an *m/z* range of 50–1,000. For each sample, 5 μL was injected twice consecutively to generate technical replicates, which were averaged prior to downstream analysis. Samples were acquired in randomized order within plates, and pooled study samples were injected regularly throughout the batch. Raw mass spectrometry data were centroided, merged, and recalibrated using algorithms adapted from ^38^. Putative metabolite annotations were generated based on accurate mass matching within a 1 mDa threshold and isotopic correlation patterns against the Human Metabolome Database (HMDB), KEGG, and ChEBI databases. Ion intensities are reported as semi-quantitative counts and are comparable across samples but not across different ions. Principal component analysis and additional multivariate analyses were performed using General Metabolics’ FiaStudio platform. Technical quality metrics, including total ion current, median ion intensity, centroid counts, and mass accuracy, were assessed using pooled study samples and were within expected bounds for the analytical run. Metabolomics data were visualized using principal component analysis (PCA) for global pattern assessment and volcano plots to display differential metabolite abundance, as indicated in the figure legends.

### Flow cytometry

Peripheral blood (100 µL per mouse) was collected at day 7 into anticoagulant-containing tubes. Red blood cells were lysed using eBioscience™ 10× RBC Lysis Buffer (Multi-species) (Thermo Fisher Scientific, Cat. no. 00-4300-54), after which leukocytes were washed and resuspended in staining buffer. Fc receptors were blocked using anti-CD16/32 antibody (clone 93 or 2.4G2), and dead cells were excluded using Zombie NIR™ Fixable Viability Dye (BioLegend, Cat. no. 423101). For surface immunophenotyping, cells were stained with fluorochrome-conjugated antibodies against CD45 (AF700), CD11b (SBUV575), Ly6C (APC-Fire 810), and Ly6G (BV785). During staining, Brilliant Stain Buffer (BD Biosciences, Cat. no. 563794) was used according to the manufacturer’s recommendations. For intracellular cytokine and transcription factor analyses, cells were stimulated *ex vivo* using eBioscience™ Cell Stimulation Cocktail (plus protein transport inhibitors), 500× (Thermo Fisher Scientific, Cat. no. 00-4975-03), as indicated. Cells were subsequently fixed and permeabilized using the eBioscience™ Foxp3 / Transcription Factor Staining Buffer Set (Thermo Fisher Scientific, Cat. no. 00-5523-00). Intracellular staining was performed using antibodies against IL-10 (APC, clone JES5-16E3; Thermo Fisher Scientific, Cat. no. 17-7101-81). Data were acquired at the UCLA Flow Cytometry Core Facility on a Sony ID7000 spectral flow cytometer and analyzed using FlowJo software (v10). Gating was performed sequentially on singlets, live cells, and CD45⁺ leukocytes, followed by subdivision into myeloid (CD11b^pos^) and lymphoid (CD11b^neg^) populations. Myeloid subsets including inflammatory monocytes, MDMCs and, neutrophils, and were identified based on differential expression of Ly6C and Ly6G. Cytokine-positive populations were quantified following stimulation, as indicated. The complete gating strategy is shown in **Supplementary Figure S3**.

### Anti–IL-10 neutralization in vivo

To block IL-10 signaling during the exposure phase, mice were treated with an IL-10–neutralizing monoclonal antibody (InVivoMAb anti-mouse IL-10, Bio X Cell, Cat. no. BE0049) or the corresponding isotype control (InVivoMAb rat IgG1 isotype control, anti-horseradish peroxidase, Bio X Cell, Cat. no. BE0088). Antibodies were administered at a dose of 200 µg per mouse.

### Statistical Analysis

All statistical analyses were performed using GraphPad Prism version 10.0.0 (GraphPad Software). Outliers were identified within Prism using a robust regression–based outlier detection method (ROUT) with a false discovery rate of 5% and were excluded from analysis. Data are presented as mean ± standard deviation (SD). Sample sizes and statistical tests are indicated in the figure legends. Longitudinal data were analyzed by two-way ANOVA and statistical significance indicates the main effect of exposure (column factor) across the time course. Fisher’s least significant difference (LSD) test was used for post hoc pairwise comparisons. When stated, individual time-point comparisons were performed using unpaired two-tailed Student’s *t* tests. A *P* value < 0.05 was considered statistically significant. Significance is denoted as ns, *P* < 0.05, \**P* < 0.01, \*\**P* < 0.001, and \*\*\**P* < 0.0001.

---

**Supplementary Figure S1.**
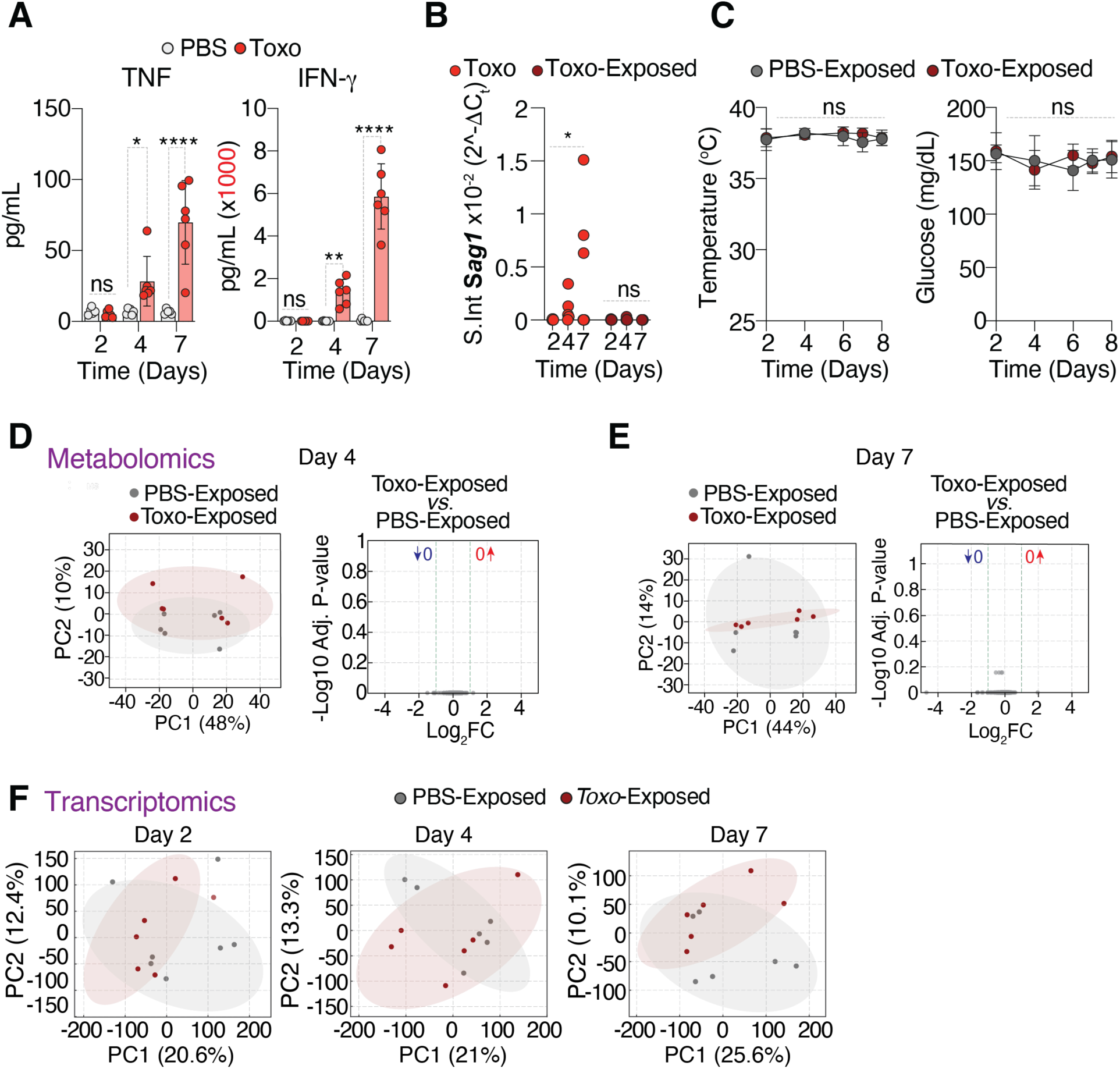
Related to Figure 1. **(A)** Serum concentrations of TNF and IL-10 measured at days 2, 4, and 7 in PBS-treated and *Toxoplasma gondii*–infected animals. Bars represent mean ± SD. Statistical analysis was performed using two-way ANOVA followed by Fisher’s LSD for pairwise comparisons. ns, not significant; \**P* < 0.05; \*\**P* < 0.01; \*\*\*\**P* < 0.0001. **(B)** Liver *Sag1* transcript abundance measured at days 2, 4, and 7 in infected (*Toxo*) and exposed (*Toxo*-exposed) animals. Data are mean ± SD. Statistical comparisons shown are day 2 versus day 7 within *Toxo* and within *Toxo*-exposed groups. ns, not significant; \**P* < 0.05 by uncorrected Fisher’s LSD (*n* = 6 mice per group). **(C)** Core body temperature and blood glucose levels of PBS-exposed and *Toxo*-exposed mice measured longitudinally throughout the exposure period. Data are mean ± SD. ns by two-way ANOVA. (*n* = 6 mice per group). **(D-E)** Principal component analysis (PCA) and volcano plot showing differential metabolite abundance between *Toxo*-exposed and PBS-exposed groups of serum metabolomics profiles from PBS-exposed and *Toxo*-exposed mice at day 4 **(D)** and 7 **(E)**. Each point represents one animal. **(F)** Principal component analysis (PCA) showing transcript abundance between *Toxo*-exposed and PBS-exposed groups of spleens from PBS-exposed and *Toxo*-exposed mice at day 2, 4 and 7. Each point represents one animal.

**Supplementary Figure S2.**
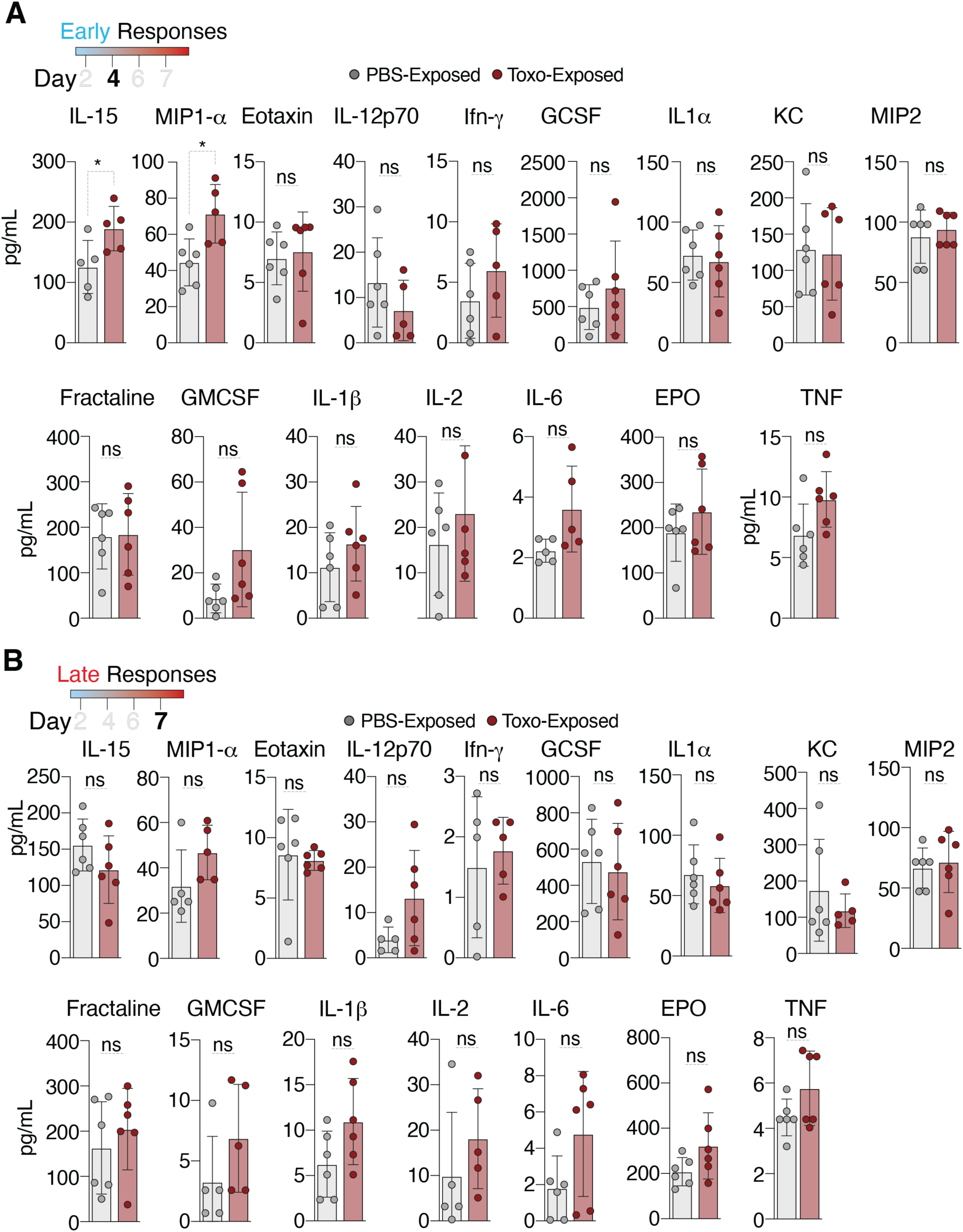
Related to Figure 2. **(A)** Serum cytokine concentrations of 4 day-PBS-exposed and *Toxo*-exposed mice. Bars represent mean ± SD. ns, not significant; \**P* < 0.05 by unpaired *t*-test. **(B)** Serum cytokine concentrations measured at day 7 in PBS-exposed and *Toxo*-exposed mice for the same cytokines shown in (A). Bars represent mean ± SD. Statistical analysis was performed using unpaired t test. ns, not significant; \**P* < 0.05 by unpaired *t*-test (*n* = 4-6 mice per group).

**Supplementary Figure S3.**
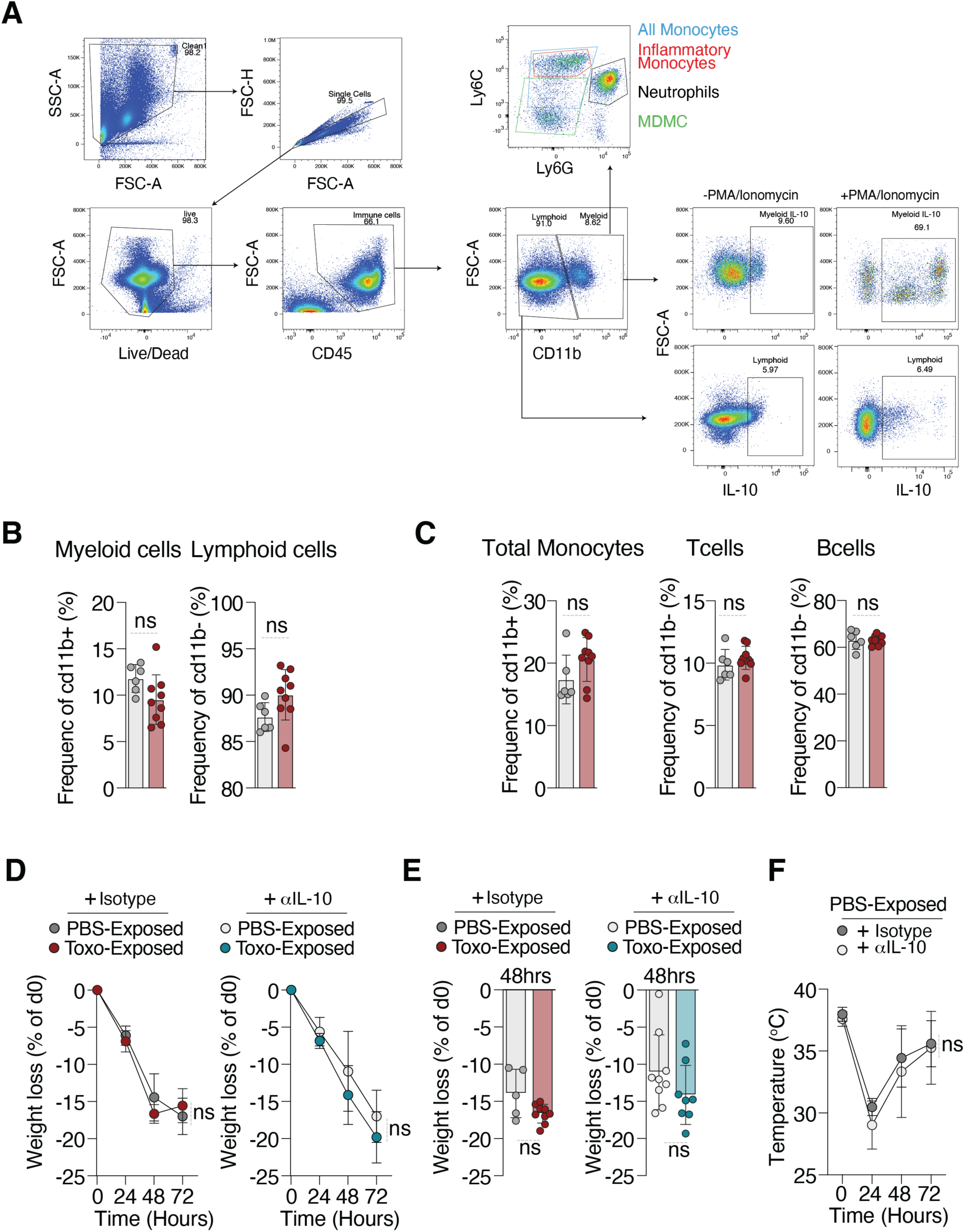
Related to Figure 3. **(A)** Flow cytometry gating strategy for identification of myeloid and lymphoid populations and IL-10 producing cells. Cells were sequentially gated on singlets, live cells, CD45⁺ immune cells, and CD11b⁺ myeloid or CD11b⁻ lymphoid populations. Myeloid subsets (including, inflammatory monocytes, monocyte-derived macrophage-like cells (MDMCs), and neutrophils, were identified based on Ly6C and Ly6G expression. Representative IL-10 staining is shown in unstimulated and PMA/ionomycin-stimulated conditions. **(B)** Frequency of CD11b^pos^ myeloid cells and CD11b^neg^ lymphoid cells in blood from PBS-exposed and *Toxo*-exposed mice. n=6 PBS-exposed mice, n=9 *Toxo-*exposed mice from two-independent experiments, data are mean ± SD. ns, not significant by unpaired *t* test. **(C)** Frequency of total monocytes, T-cells and B cells in blood from PBS-exposed and *Toxo*-exposed mice. n=6 PBS-exposed mice, n=9 *Toxo-*exposed mice from two-independent experiments, data are mean ± SD; ns, not significant by unpaired *t* test. B cells were gated as CD11b⁻Ly6G⁻Ly6C⁻CD3⁻CD19⁺MHCII⁺, and T cells as CD11b⁻Ly6G⁻Ly6C⁻CD3⁺CD19⁻. **(D)** Body weight loss at indicated hours post LPS challenge in 6-day PBS-exposed and *Toxo*-exposed mice treated with isotype control (left) or anti-IL-10 antibody (right). Line plots show longitudinal measurements. Data are mean ± SD from mice as in (B) by two-way ANOVA analysis. **(E)** Body weight loss at 24 and 48 hours post LPS challenge in mice as in (D). Data are mean ± SD. and individual time-point comparisons were performed by unpaired t test. ns, not significant. **(F)** Core body temperature trajectories following LPS challenge in PBS-exposed mice treated with isotype control or anti-IL-10 antibody at indicated time points. Data are mean ± SD from mice as in (B) by two-way ANOVA analysis.

**Figure S4:**
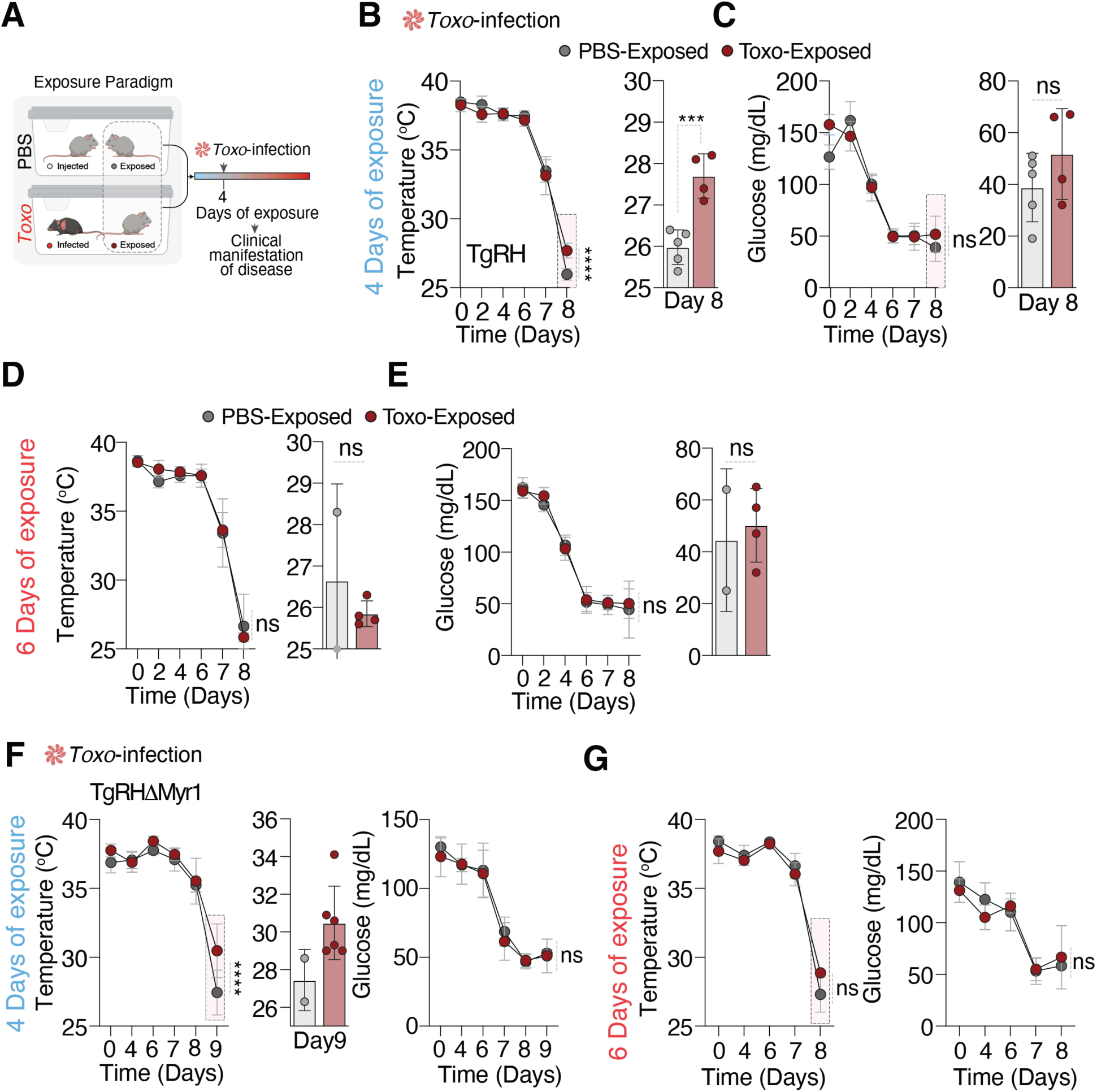
Exposure to infected cage-mates promotes thermoregulatory resilience extends during live *Toxoplasma gondii* infection. **(A)** Schematic of the exposure paradigm. Mice were exposed to PBS-treated or *Toxo*-infected cage mates for 4 or 6 days, followed by infection with the virulent *T. gondii* RH strain or a less virulent *Toxoplasma gondii* RHΔ. Disease progression was monitored until mice had to be euthanized in accordance with humane end points. **(B)** Core body temperature during live *Toxo*-infection in mice exposed for 4 days. Line plots show longitudinal measurements; bar plot shows temperature at day 8. Data are mean ± SD. Trajectories were analyzed by two-way ANOVA; pairwise comparison at day 8 was performed by unpaired *t* test; \*\*\**P* < 0.001; \*\*\*\**P* < 0.0001. Starting sample size n = 6 mice per group. At day 8, analyzed mice (*n* = 5 mice in PBS-exposed; 4 mice in *Toxo*-exposed) were mice whose morbidity were in accordance with animal welfare protocols while the other mice were euthanized in accordance with humane end point. **(C)** Blood glucose levels during live *T. gondii* infection in mice exposed for 4 days. Line plots show longitudinal measurements; bar plot shows glucose at day 8. Data are mean ± SD. Trajectories were analyzed by two-way ANOVA; pairwise comparison at day 8 was performed by unpaired *t* test. ns, not significant. Similar sample size consideration as in B. **(D)** Core body temperature and **(E)** blood glucose, during live *T. gondii* infection in mice exposed for 6 days. Data are shown as longitudinal measurements and bar plot shows temperature at day. Mean ± SD. Trajectories were analyzed by two-way ANOVA. ns, not significant. All data are from one experiment. Similar sample size consideration as in B. **(F)** Core body temperature and blood glucose during live infection with *Toxoplasma gondii* RHΔmyr1 in mice exposed for 4 days. Line plots show longitudinal measurements; bar plot shows temperature at day 9. Data are mean ± SD. Trajectories were analyzed by two-way ANOVA, \*\*\*\**P* < 0.01; statistical pairwise comparison was not conducted due to *n* = 2 at day 9 for PBS-exposed. Similar sample size consideration as in B, i.e starting sample size *n* = 6. **(G)** Core body temperature and blood glucose during live infection with *Toxoplasma gondii* RHΔmyr1 in mice exposed for 6 days. Line plots show longitudinal measurements; bar plot shows temperature at day 9. Data are mean ± SD. Trajectories were analyzed by two-way ANOVA; n = 3 mice per group. All data from one experiment.

**Table S1.** Day 4 cytokines in PBS-exposed vs *Toxo*-exposed mice. Serum cytokine concentrations at day 4 are shown as mean ± SD (n=6 mice per group). P values were calculated by two-tailed unpaired t-test (PBS-exposed vs *Toxo*-exposed) for each cytokine.

| Cytokine | PBS-exposed mean (± SD) | Toxo-exposed mean (± SD) | p-value (unpaired t-test) |
| --- | --- | --- | --- |
| IL-3 | 0.22 (± 0.11) | 0.23 (± 0.10) | 0.84 |
| IL-7 | 4.63 (± 3.15) | 5.16 (± 3.84) | 0.81 |
| IL-9 | 3.01 (± 0.00) | 3.01 (± 0.00) | 1 |
| IL-12p40 | 29.87 (± 11.86) | 26.21 (± 14.08) | 0.64 |
| IL-13 | 25.31 (± 12.34) | 44.48 (± 27.94) | 0.16 |
| IL-17 | 1.57 (± 0.37) | 2.51 (± 1.19) | 0.09 |
| IP-10 | 92.47 (± 15.71) | 112.12 (± 33.38) | 0.22 |
| LIF | 1.81 (± 0.84) | 1.53 (± 0.21) | 0.55 |
| LIX | 1070.61 (± 884.78) | 651.72 (± 408.78) | 0.32 |
| M-CSF | 5.29 (± 1.28) | 7.87 (± 4.49) | 0.21 |
| MCP-1 | 62.87 (± 31.22) | 65.68 (± 31.26) | 0.88 |
| MIG | 3461.20 (± 1328.18) | 4135.18 (± 1716.01) | 0.46 |
| MIP-1β | 47.31 (± 17.38) | 118.12 (± 83.16) | 0.07 |
| RANTES | 29.83 (± 13.19) | 33.56 (± 9.92) | 0.59 |
| VEGF | 0.54 (± 0.33) | 1.15 (± 0.82) | 0.13 |
| IFNβ-1 | 190.60 (± 106.25) | 338.24 (± 247.43) | 0.21 |
| IL-16 | 2647.01 (± 976.57) | 1721.15 (± 1170.74) | 0.17 |
| IL-20 | 90.78 (± 53.05) | 90.96 (± 27.37) | 0.99 |
| MCP-5 | 616.17 (± 617.69) | 328.82 (± 169.92) | 0.30 |
| MDC | 291.72 (± 97.52) | 252.65 (± 78.69) | 0.46 |
| MIP-3α | 25.53 (± 4.76) | 30.83 (± 6.47) | 0.14 |
| MIP-3β | 92.19 (± 33.26) | 94.61 (± 28.81) | 0.90 |
| TARC | 630.97 (± 282.05) | 611.79 (± 289.64) | 0.91 |

**Table S2.** Day 7 cytokines in PBS-exposed vs *Toxo*-exposed mice. Serum cytokine concentrations at day 7 are shown as mean ± SD (n=6 mice per group). P values were calculated by two-tailed unpaired t-test (PBS-exposed vs *Toxo*-exposed) for each cytokine. Bold case identifies significantly different cytokine.

| Cytokine | PBS-exposed mean (± SD) | Toxo-exposed mean (± SD) | p-value (unpaired t-test) |
| --- | --- | --- | --- |
| IL-3 | 0.20 (± 0.11) | 0.20 (± 0.11) | 1.00 |
| IL-7 | 2.06 (± 1.61) | 3.70 (± 2.85) | 0.25 |
| IL-9 | 5.00 (± 3.16) | 8.77 (± 14.12) | 0.54 |
| IL-12p40 | 18.29 (± 7.38) | 21.59 (± 6.61) | 0.43 |
| IL-13 | 23.12 (± 6.01) | 29.04 (± 24.01) | 0.61 |
| IL-17 | 0.67 (± 0.46) | 3.52 (± 2.75) | **0.03** |
| IP-10 | 88.81 (± 27.47) | 100.85 (± 20.27) | 0.41 |
| LIF | 1.35 (± 0.37) | 1.41 (± 0.68) | 0.85 |
| LIX | 943.12 (± 582.39) | 180.69 (± 26.57) | **0.03** |
| M-CSF | 4.08 (± 0.82) | 4.08 (± 1.24) | 1.00 |
| MCP-1 | 36.59 (± 17.60) | 37.38 (± 20.48) | 0.94 |
| MIG | 3084.77 (± 818.14) | 3514.07 (± 947.69) | 0.42 |
| MIP-1β | 40.74 (± 7.54) | 61.70 (± 29.89) | 0.16 |
| RANTES | 29.10 (± 11.78) | 27.59 (± 4.46) | 0.77 |
| VEGF | 0.37 (± 0.24) | 0.50 (± 0.22) | 0.39 |
| IFNβ-1 | 119.38 (± 35.18) | 257.03 (± 151.20) | **0.05** |
| IL-16 | 2149.29 (± 1246.99) | 2667.20 (± 934.12) | 0.43 |
| IL-20 | 51.94 (± 20.95) | 59.96 (± 15.37) | 0.47 |
| MCP-5 | 243.60 (± 162.16) | 554.29 (± 228.61) | **0.02** |
| MDC | 219.12 (± 154.46) | 216.53 (± 83.07) | 0.97 |
| MIP-3α | 25.49 (± 13.60) | 24.85 (± 7.88) | 0.92 |
| MIP-3β | 58.44 (± 11.90) | 87.75 (± 27.94) | **0.04** |
| TARC | 607.75 (± 353.85) | 908.80 (± 700.38) | 0.37 |

